# Automated detection of artefacts in neonatal EEG with residual neural networks

**DOI:** 10.1101/2021.05.23.445349

**Authors:** Lachlan Webb, Minna Kauppila, James A. Roberts, Sampsa Vanhatalo, Nathan Stevenson

## Abstract

**Background and Objective:** To develop a computational algorithm that detects and identifies different artefact types in neonatal electroencephalography (EEG) signals.

**Methods:** As part of a larger algorithm, we trained a Residual Deep Neural Network on expert human annotations of EEG recordings from 79 term infants recorded in a neonatal intensive care unit (112 h of 18-channel recording). The network was trained using 10 fold cross validation in Matlab. Artefact types included: device interference, EMG, movement, electrode pop, and non-cortical biological rhythms. Performance was assessed by prediction statistics and further validated on a separate independent dataset of 13 term infants (143 h of 3-channel recording). EEG pre-processing steps, and other post-processing steps such as averaging probability over a temporal window, were also included in the algorithm.

**Results:** The Residual Deep Neural Network showed high accuracy (95%) when distinguishing periods of clean, artefact-free EEG from any kind of artefact, with a median accuracy for individual patient of 91% (IQR: 81%-96%). The accuracy in identifying the five different types of artefacts ranged from 57%-92%, with electrode pop being the hardest to detect and EMG being the easiest. This reflected the proportion of artefact available in the training dataset. Misclassification as clean was low for each artefact type, ranging from 1%-11%. The detection accuracy was lower on the validation set (87%). We used the algorithm to show that EEG channels located near the vertex were the least susceptible to artefact.

**Conclusion:** Artefacts can be accurately and reliably identified in the neonatal EEG using a deep learning algorithm. Artefact detection algorithms can provide continuous bedside quality assessment and support EEG review by clinicians or analysis algorithms.

**Highlights:**

- We applied a Residual Deep Neural Network as part of an artefact detection algorithm in neonatal electroencephalograms.
- The algorithm shows high accuracy in identifying artefactual data in general and for specific artefact types.
- EEG channels near the top of the head are less prone to artefact.

## 1. Introduction

Monitoring with electroencephalography (EEG) is a standard of care in many neonatal intensive care units (NICU) [1]. It is frequently used to detect neurological abnormalities, evaluate maturation, and predict neurodevelopmental outcome [2-10]. The rapid uptake of long term EEG monitoring in the NICU has uncovered two, as yet unresolved bottlenecks: how to record reliable, high quality EEG, and how to provide real-time interpretation for treating clinicians [11]. In this study, we focus on the problem of ensuring that a reliable, high quality EEG signal is recorded for clinical review [3, 12-14]. Recording reliable EEG requires real-time surveillance of EEG signal quality. This is traditionally achieved by visual signal inspection at the cotside. However, it requires constant attention and substantial expertise in EEG reading, neither of which can be expected from nursing staff whose chief responsibility is immediate care of the patient. The only practically feasible solution would be to develop an automated measure of signal quality that allows real time feedback to the nursing staff. In practise, such an automated tool will identify the type of artefact so that the nursing staff can troubleshoot the signal quality issues.

Several methods for have been developed for the detection [15, 16] and removal [17-19] of artefacts in adult EEG. These methods have limited applicability to neonatal EEG due to: i) the difference in manifestation, variability and susceptibility of artefacts, ii) fundamental differences in predominant EEG patterns, and iii) a lower number of recording electrodes. To overcome these challenges, researchers have attempted to develop artefact detection methods specific to neonatal EEG, but these methods have used i) small datasets, ii) infants without pathology, and iii) not accurately classified the type of artefact [5, 14, 20, 21]. Recent advances in machine learning (notably, deep neural networks [22]) provide an opportunity to improve the detection performance towards clinical applicability if sufficiently large datasets of annotated neonatal EEG can be acquired.

Here, we present an algorithm for detecting artefacts in neonatal EEG using a deep residual neural network. The performance of this algorithm was evaluated on datasets representative of data typically obtained in a NICU setting. We also validated the algorithm on an independent dataset.

## 2. Material and Methods

### 2.1 Data acquisition

#### 2.1.1 EEG Recordings

This study consists of continuous multichannel EEG recordings from 79 term neonates with the average gestational age (GA) of 40 weeks (range 35-44 weeks). The recordings were performed in the neonatal intensive care unit in the Hospital District of Helsinki and Uusimaa, and the Helsinki University Central Hospital (HUCH) specialist medical care area. EEGs were recorded using the NicoletOne vEEG system with sampling frequency of 256 Hz. A total of 21 electrodes were placed according to the international 10-20 system using a referential montage. A bipolar montage with 18 channels (Fp2-F8, F8-T4, T4-T6, T6-O2, Fp1-F7, F7-T3, T3-T5, T5-O1, Fp2-F4, F4-C4, C4-P4, P4-O2, Fp1-F3, F3-C3, C3-P3, P3-O1, Fz-Cz, Cz-Pz) was used for analysis. The average length of the recordings for each patient was 85 min (range 52-257 min). This dataset was initially compiled to evaluate seizure detection by human experts and computer algorithms and is publicly available [23, 24]. These infants represent a typical cohort that would be monitored in the NICU.

A validation dataset (n**=**13) was collected from a series of infants monitored for seizures with subsequent detection of stroke in a clinical database of long-term monitoring (Helsinki University Central Hospital, Finland). Elements of this patient cohort have been described and published earlier [25]. EEGs were recorded using frontal and parietal electrodes (F3, F4, P3, P4) with a NicoletOne vEEG system (same recording device) resampled to a frequency of 256 Hz (median recording duration 9 hours, IQR: 8 hours to 16 hours).

#### 2.1.2 Data Annotation

A human reviewer (NJS) initially identified obvious periods of clean artefact-free EEG and several types of artefacts, annotating the start and end times. An experienced clinical neurophysiologist (SV) then reviewed these annotations and accepted, modified, or deleted each accordingly (5% disagreement on identified events). A final reviewer (LW) selected the specific EEG derivations that presented with artefact or clean EEG, i.e. assigned the event annotation to the specific channels. Artefact classes identified included: *Device Interference* artefacts (*DI*; interfering electromagnetic radiation from nearby devices), *Electromyography* artefact (*EMG*; muscle activation), *Movement* (*MO*; long, sustained movement of the head), *Electrode Pop* (*EL*; transients due to poor electrode contact), and repetitive *Biological Rhythm* artefact (*BIO*; cardiac or respiratory origin) [6]. Examples are shown in Figure 1.

**Figure 1:**
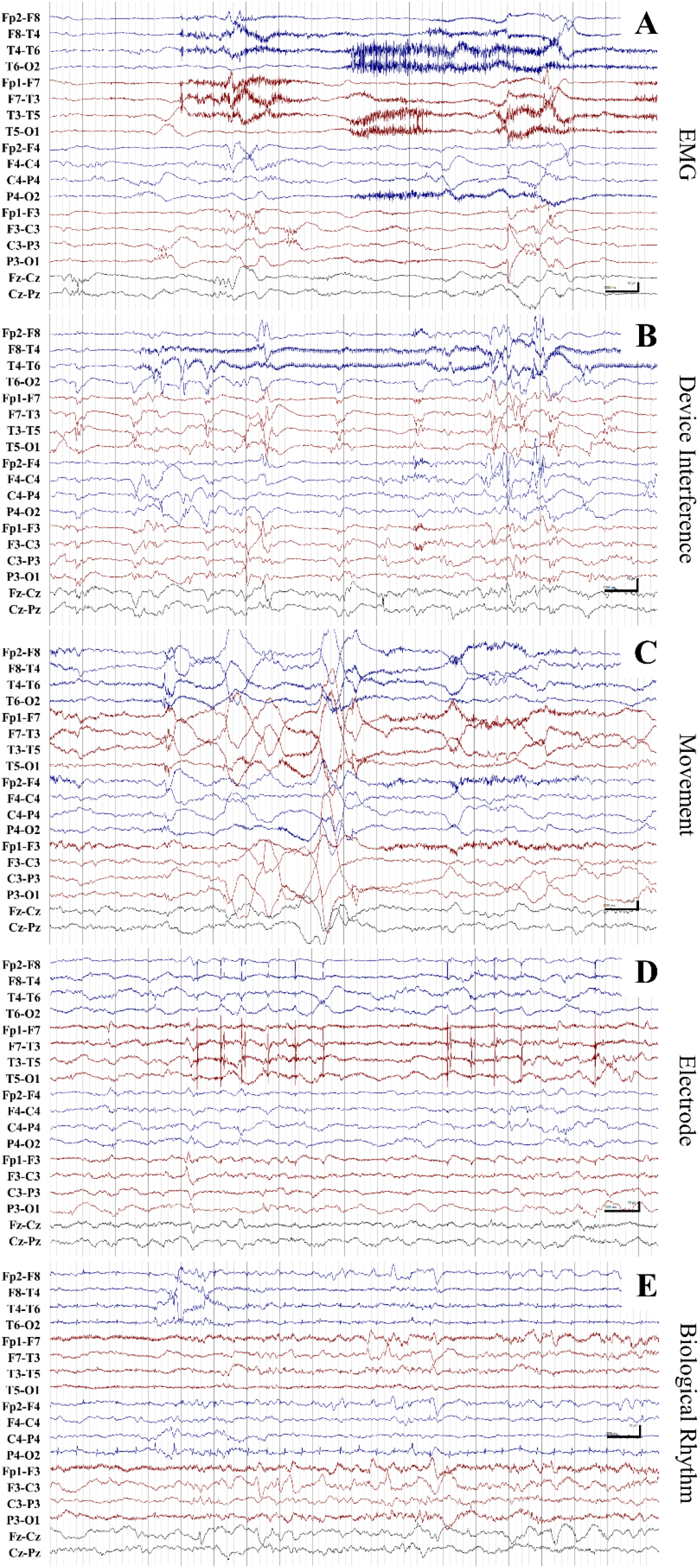
Exemplar segments of EEG recording for different types of artefact. A) Muscle artefact on predominantly on the temporal regions (*EMG*). B) *Device Interference* on T4. C) *Movement* artefact on F8, F7, T3, and P3 (muscle artefact is also present). D) *Electrode pop* artefact predominantly on T3. E) ECG artefact (example of the *Biological Rhythm* class) on P4-O2. The calibration scale markers denote 1 s horizontally and 70 µV vertically.

Out of the 112 hours of 18-channel EEG recordings (2014 hours of single channel recordings), 44 hours of annotated events (Supplementary Table S.1) resulted in 277 hours of single-channel EEG data, distributed across 6 classes.

The annotated periods of EEG were then further divided into segments of 4 seconds in duration with 50% (2 second) overlap (See Table 1, Unbalanced). Given the imbalance in the prevalence of each class of event, and a balanced dataset being more conducive to training the deep neural network, different levels of overlap were used for each class to achieve a more balanced dataset for training. Clean EEG was the most common class, and so not only had no overlap, but a random half of the data were also deleted. The EMG segments had 2 second (50%) overlap, and *MO* artefact segments had 3 second (75%) overlap. *EL* events had a 3.5 second (87.5%) overlap, and the remaining artefacts (*BIO* and *DI*) had a 3.75 second (93.75%) overlap.

**Table 1:** Segment descriptive statistics. The Unbalanced Data has all segments with 50% overlap. The Balanced Data has artefact class specific overlap and half of the Clean segments removed.

| Labelled signal | Unbalanced |  |  | Balanced |
| --- | --- | --- | --- | --- |
|  | Number of 4 sec segments (%) | Patients with event (n) | Avg. number segments, median (IQR)^ | Number of 4 sec segments (%) |
| Clean EEG | 337130 (69.8%) | 79 | 3458 (2059.3 – 5386) | 84760 (21.0%) |
| Device Interference (DI) | 8601 (1.8%) | 16 | 190.5 (18 – 572) | 62302 (15.4%) |
| Electromyography (EMG) | 80361 (16.6%) | 74 | 708 (215 – 1469) | 80361 (19.9%) |
| Movement (MO) | 35452 (7.3%) | 52 | 210.5 (85.5 – 718.5) | 68893 (17.1%) |
| Electrode Pop (EL) | 14099 (2.9%) | 73 | 54 (26.7 – 171) | 48434 (12.0%) |
| Biological Rhythms (BIO) | 7493 (1.6%) | 14 | 227.5 (119– 952) | 59044 (14.6%) |

^ average number in patients with event type

For the validation data, a section of recording that contained several types of artefact was extracted from each EEG. The section was continuously annotated (F3-P3, F4-P4, P3-P4) for artefacts and periods of artefact-free EEG by an experienced clinical neurophysiologist (SV). The type of artefacts to be annotated were limited to the 5 classes used in the training dataset. This process of annotation of the validation dataset was different to that used when acquiring the training dataset. While the trained dataset extracted obvious, exemplar segments of artefact and clean EEG from recordings, the continuous annotation of the validation dataset required increased decision making relating to less obvious artefacts resulting in increased ambiguity.

### 2.2 Artefact detection algorithm

The artefact detection algorithm input was a 4 s EEG segment and the output was a class label: *Clean* uncontaminated EEG, *DI, EMG, MO, EL*, and *BIO*. The algorithm consisted of a pre-processing of raw EEG data stage, a neural network, and a post-processing stage (Figure 2).

**Figure 2:**
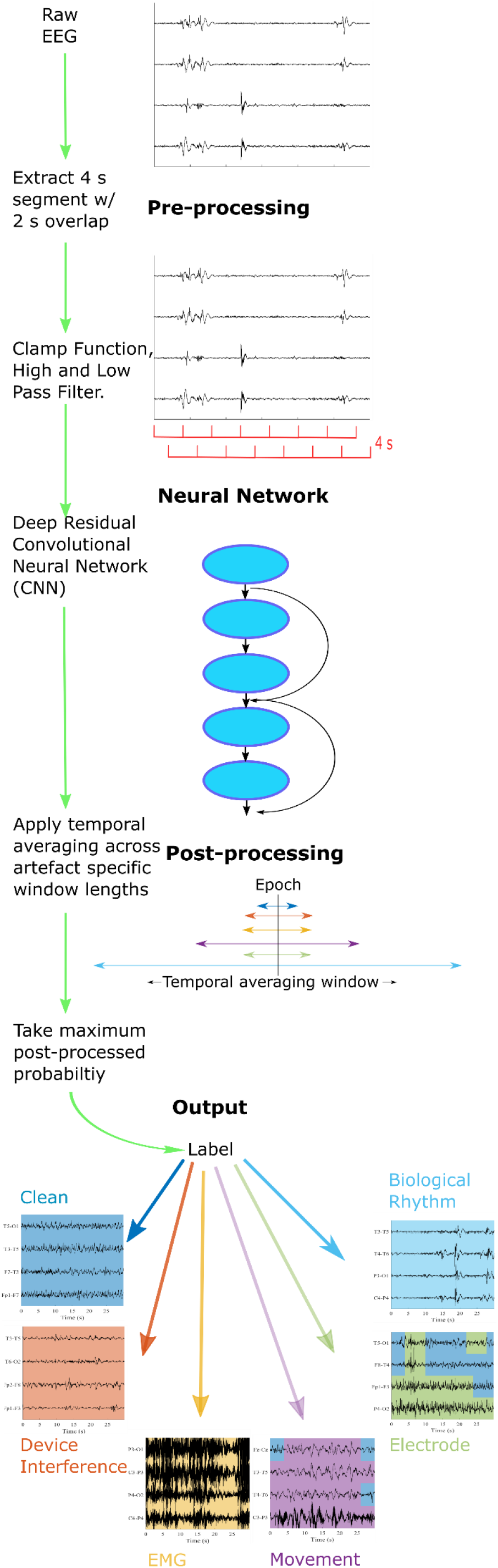
Flow diagram of artefact detection system. Green arrows denote the algorithm implementation of data pre-processing, the deep residual network, and post-processing.

Our pre-processing steps took the 4 s segments of EEG data and applied a clamping function (Equation 1) to limit the dynamic amplitude range of the segments, yielding clamped segments *EEG*_*c*_(*t*) given by

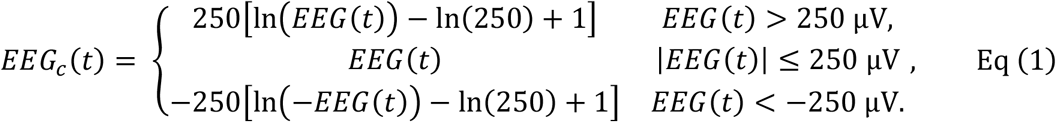

These EEG segments were then filtered with a Butterworth high pass filter (2-poles, cut-off frequency of 0.5 Hz) and a 50 Hz notch filter (2-pole, IIR filter). Each EEG segment was then processed by a neural network. Several network architectures were trialled (see Section 2.3).

In our post-processing stage, the output of the softmax layer (a vector the length of the number of event classes representing the probability of the processed segment belonging to a particular class) was averaged over a time period specific to each artefact type (temporal smoothing) to take into account the fact that nearby segments in time are more likely to have the same label and some artefact types present with different durations. The artefact class with the highest smoothed probability was chosen as the final artefact label [26].

### 2.3 Training, testing and evaluation of the artefact detector

Two different neural network architectures were considered, a deep residual network [27] and a classical deep neural network [28]. A number of depth and filter size combinations were considered for each architecture (see Table S.3). The performance of these networks was also compared to a ‘bag of features’ classified by a Support Vector Machine (SVM) [5].

Artefact detection algorithms were evaluated using 10-fold cross-validation on the Balanced Dataset. As the dataset contained EEG recordings from 79 infants, the 10-fold cross-validation used nine groups of eight patients and one of seven patients. The patients were randomly assigned to the 10 groups, i.e. all data from an individual patient is in one fold, though a seed was selected to ensure that each fold contained all event types. The number of segments used in each training fold are summarized in Supplementary material (Table S.2).

Deep neural networks were trained using the “Stochastic Gradient Descent with Momentum” algorithm (SGDM) [29]. Data was trained for 50 epochs with a Minibatch Size of 1024 and momentum set to 0.9. The learning rate was initially set at 0.1 and then reduced by a factor of 0.2 every 30 epochs. The MATLAB function trainNetwork was used to train the network in each fold.

The optimal network was selected based on a criteria of network size, binary and 6-class accuracy (see section 2.4 for definitions of accuracy). The optimal network was further tested with and without the pre-processing stage and several different training options (minibatch size, solver type, See Table S.3).

The typical event duration differs between classes, with artefacts like Electrode typically lasting 4 seconds and Biological Rhythms lasting 40 seconds (see Supplementary Table S.1). Temporal averaging was trialled by taking the average of the artefact class probabilities in surrounding segments. A greedy algorithm was used to find the optimal temporal window size for each event class on the entire Balanced Dataset. For each artefact class, several window durations (0 - 30s) were applied, while all other classes were not windowed. The window duration and class that maximized the 6 Class accuracy was then selected, and the process repeated until all artefact classes had an optimal window. The greedy algorithm was also applied separately to each fold in the Balanced Data.

### 2.4 Algorithm performance assessment

We visualized the performance of the artefact detection system by calculating the confusion matrix of the pooled data (six by six). Initial evaluation was based on the Balanced Dataset. We further summarized the performance by calculating the 6 Class accuracy, defined as the percentage of segments correctly classed as either *Clean* or correctly classed as one of the five artefact classes (Equation 2). We also evaluated the performance of the algorithm for differentiating between clean EEG and artefact (a binary decision combining all artefacts types). Formulae for the statistics considered a 4 second segment of data that was correctly identified as *Clean* as True Positive (TP), a correctly identified segment of artefact as True False (TF), a segment misclassified as *Clean* as False Positive (FP), and a segment misclassified as artefact a False Negative (FN). These binary statistics included accuracy (Equation 3), sensitivity (the ratio of true clean segments over all the segments labelled as *Clean*, Equation 4), area under the receiver operating characteristic curve (AUC, based on the probability of the *Clean* label in the softmax layer), and F1 (a measure combining sensitivity and precision (Equation 5), where precision is the ratio of segments correctly labelled as clean out of the all the segments annotated as clean, Equation 6) [30]. All measures were calculated on the pooled data (n=79).

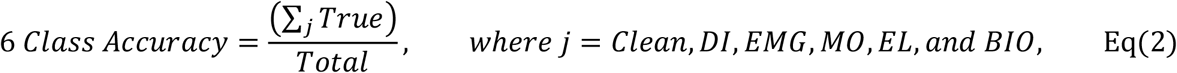

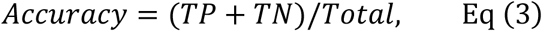

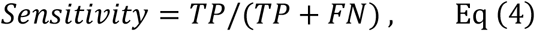

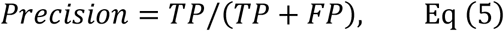

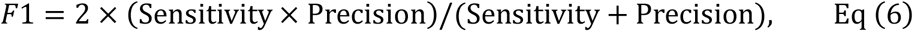

To provide a visualisation of the separation of the artefacts by the deep residual network, we took the output from a late stage of the network (fully connected layer, see Figure S.1) for 10% of the data (Balanced Data). This 6 dimensional output was reduced to 2 using UMAP [31] to visualise classifier performance. In the 2D space of the UMAP output, high performing classification manifests as clusters of similarly-identified points.

To simulate a potential application of the artefact detection algorithm (trained on the Balanced Dataset), we applied it in conjunction with a seizure detection algorithm [32] to the complete EEG recordings from all 79 patients to estimate the level of contamination within our database. The level of contamination was compared between channels in the bipolar montage.

The performance measures were also calculated for the validation dataset to show how the algorithm performed on a separate dataset. Bootstrapped confidence intervals were calculated to compare the 6-Class and binary accuracy between training and validation dataset.

## 3. Results

Results from the optimization of network architecture and training options (trained on the Balanced Dataset) were calculated from both the Balanced and Unbalanced Dataset. We tested the algorithm on the Unbalanced Dataset as this best reflects the real world clinical setting where the majority of data is typically clean and artefact types have different occurrence frequencies. We show example recordings where the algorithm correctly and incorrectly identifies artefacts, and visualize the network outputs to show the separation and overlap of the classes to support the distribution of accuracy within the confusion matrix.

We show the usefulness of the algorithm by estimating the amount of artefact present in typical EEG recordings and then validate the artefact detection algorithm on an independent, unseen dataset that was annotated under a continuous monitoring paradigm rather than the exemplar paradigm used to generate the training set. Finally, we show differences in EEG characteristics between the two annotation paradigms that help to explain the reduction in validation performance.

### 3.1 Training Results

The primary results from the Balanced Dataset show the optimal implementation of the algorithm was achieved using a residual neural network with pre-processing and temporal smoothing (Network 2 in Supplementary Figure S.2).

The optimal network was trained with SGDM and a minibatch size of 2048 (Supplementary Figure S.2). The majority of permutations of network architecture and training options gave similar results with the binary accuracy ranging from 92% to 95%, and 6-Class accuracy from 75% to 82%. Residual networks consistently outperformed networks without feed forward stages.

The binary accuracy of the artefact detection algorithm (trained on the Balance Dataset) for differentiating between artefactual EEG and clean, uncontaminated EEG was 95% and 97% when applied to the Balanced and Unbalanced Datasets, respectively. Temporal averaging improved the accuracy further with the largest improvement seen in the Balanced Dataset (Table 2). The median binary accuracy (for *Clean* vs any artefact) with temporal averaging applied across the cohort of 79 was 98% (IQR: 95% - 99%) for the Balanced Dataset and 99% (IQR: 95% - 100%) for the Unbalanced Dataset. The *Clean* EEG and *EMG* artefact were the most accurately detected (95% and 92%, Figure 5), while *DI* and *EL* artefacts were detected with the lowest accuracy (75% and 57%; Figure 5). The algorithm accuracy reflects the low proportion of *DI* and *EL* in the training dataset. While the accuracy for correctly identifying the type of artefact was low for *DI* and *EL* artefact, the binary accuracy with the true event type for detecting any type of artefact was high (97% and 91%, Figure 5). Hence, the artefacts are reliably detected, but their specific type may be identified at lower accuracy.

**Table 2:** Artefact detection performance statistics on Balanced Data and Unbalanced Data, with and without temporal smoothing. The results from the SVM [5] are also included and best compared to the Residual Neural Network trained on Balanced Data.

|  | 6 Class | Clean vs Artefact |  |  |  |
| --- | --- | --- | --- | --- | --- |
|  | Accuracy | Accuracy | Sensitivity | AUC | F1 |
| Residual Neural Network trained on Balanced Data | 82.0% | 94.5% | 95.02% | 0.947 | 0.880 |
| Residual Neural Network trained on Balanced Data and Temporal Smoothing | 84.8% | 95.5% | 97.5% | 0.962 | 0.900 |
| Residual Neural Network applied to Unbalanced Data | 91.9% | 95.4% | 95.0% | 0.957 | 0.966 |
| Residual Neural Network applied to Unbalanced Data with temporal smoothing | 94.5% | 97.2% | 97.4% | 0.971 | 0.980 |
| Bag of Features method (SVM) | 68.8% | 89.2% | 87.5% | 0.886 | 0.773 |

The optimal window lengths (in order: *Clean, DI, EMG, MO, EL*, and *BIO*) were 8, 12, 12, 28, 12, and 56 seconds which correspond to durations of 3, 5, 5, 13, 5, and 27 segments respectively. These values were supported by the results from applying the greedy algorithm to each fold separately (Table S.4). These window lengths align with the median length of the annotated artefact events (Table S.2), with *MO* and *BIO* typically being longer in duration. Example outputs of the algorithm superimposed over a period of EEG recording are shown in Figures 3 and 4.

**Figure 3.**
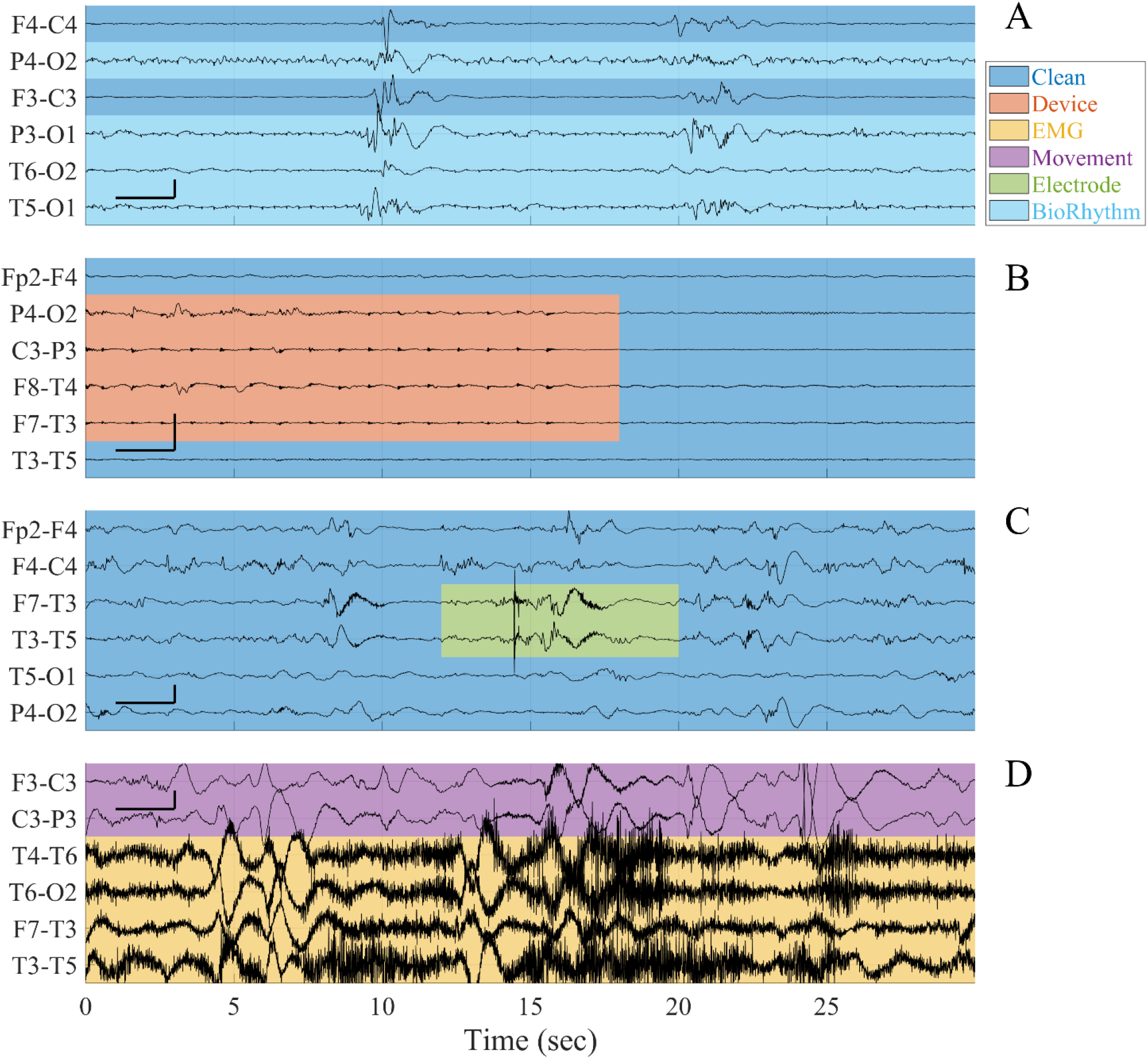
Examples of accurately detected artefacts. Colours denote the algorithm classifications. A) *Biological Rhythm* correctly identified. Annotation was on P4-O2, P3-O1, T6-O2, and T5-O1, with the clearest annotation on T5-O1 for the entire segment. B) *Device Interference* correctly identified. The annotation was on P4-O2, C3-P3, F8-T4, F7-T3 (0-18s). C) *Electrode pop* correctly identified. *Electrode pop* annotated on F7-T3 and T3-T5 (13-20s). D) *Movement* and *EMG* correctly identified. *EMG* annotated on T4-T6, T6-O2, F7-T3, T3-T5 (entire segment). *Movement* annotated on F3-C3 and C3-P3 (entire segment). The calibration scale bars denote 2 s horizontally and 100 µV vertically.

**Figure 4.**
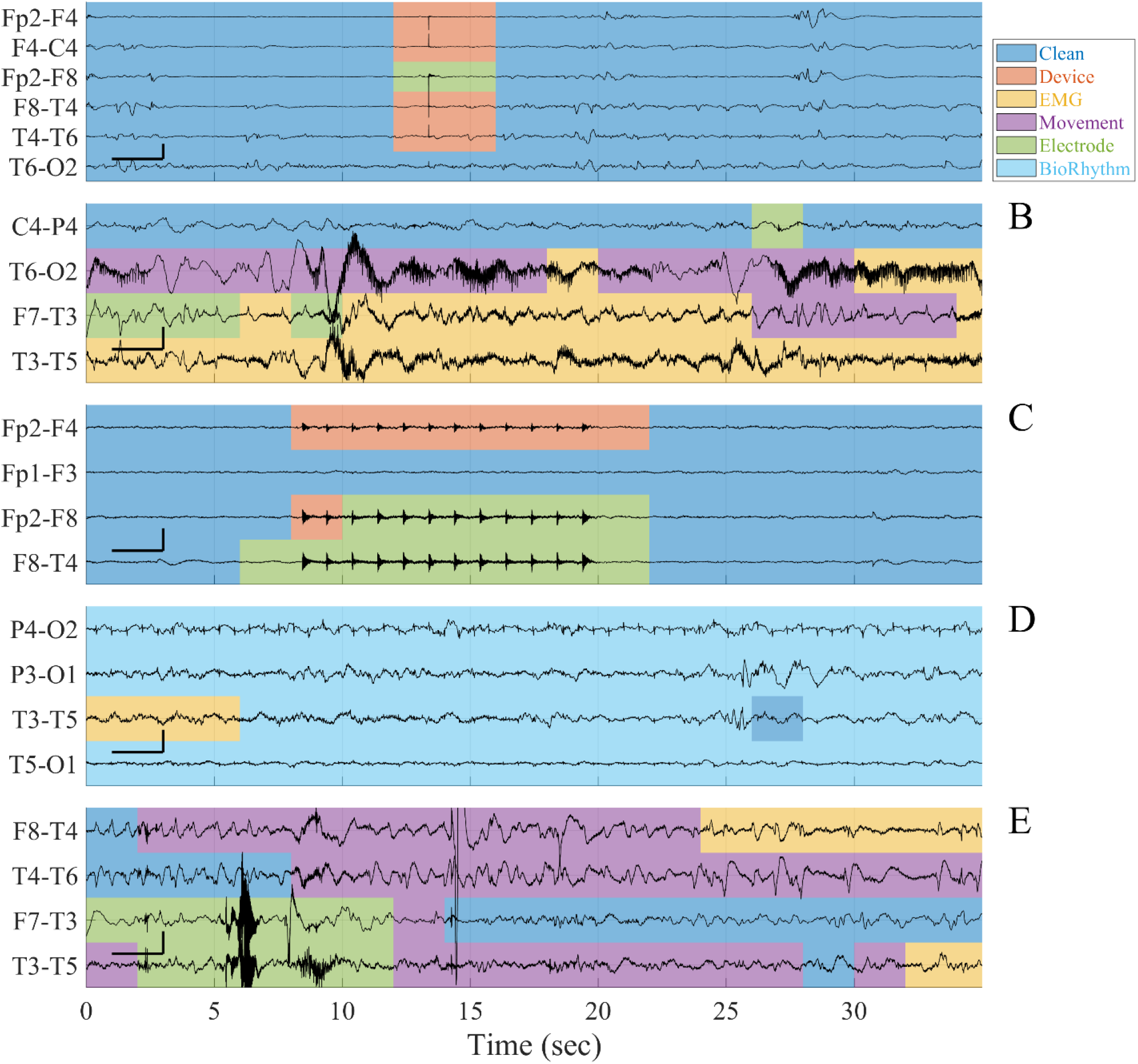
Examples of algorithm misclassifications. Colours denote the algorithm classifications. A) *Electrode* annotated on Fp2-F4, F4-C4, Fp2-F8, F8-T4, T4-T6, T6-O2 (12 to 16 s) was labelled by the algorithm as *Device Interference* on Fp2-F4, F4-C4, F8-T4, T4-T6. The *Electrode* artefact on Fp2-F8 was correctly identified. B) *EMG* annotated on T6-O2 (8 to 22 s) was labelled by the algorithm as *Movement. Movement* annotated on F7-T3 and T3-T5 (0 to 30 s) was labelled by the algorithm as *Electrode* and *EMG*. C) *Device Interference* was annotated on Fp2-F4, Fp2-F8, F8-T4 (8 to 20 s) was labelled as *Electrode* artefact on Fp2-F8, F8-T4 and correctly identified on Fp2-F4. D) *Biological Rhythm* annotated on P4-O2, P3-O1, T3-T5, T5-O1 (entire segment) was labelled, in part, by the algorithm as *EMG* and *Clean* on T3-T5. E) *Electrode* annotated on F8-T4, T4-T6 (14 to 18 s) was labelled as *Movement*. The calibration scale bars denote 2 s horizontally and 100 µV vertically.

**Figure 5:**
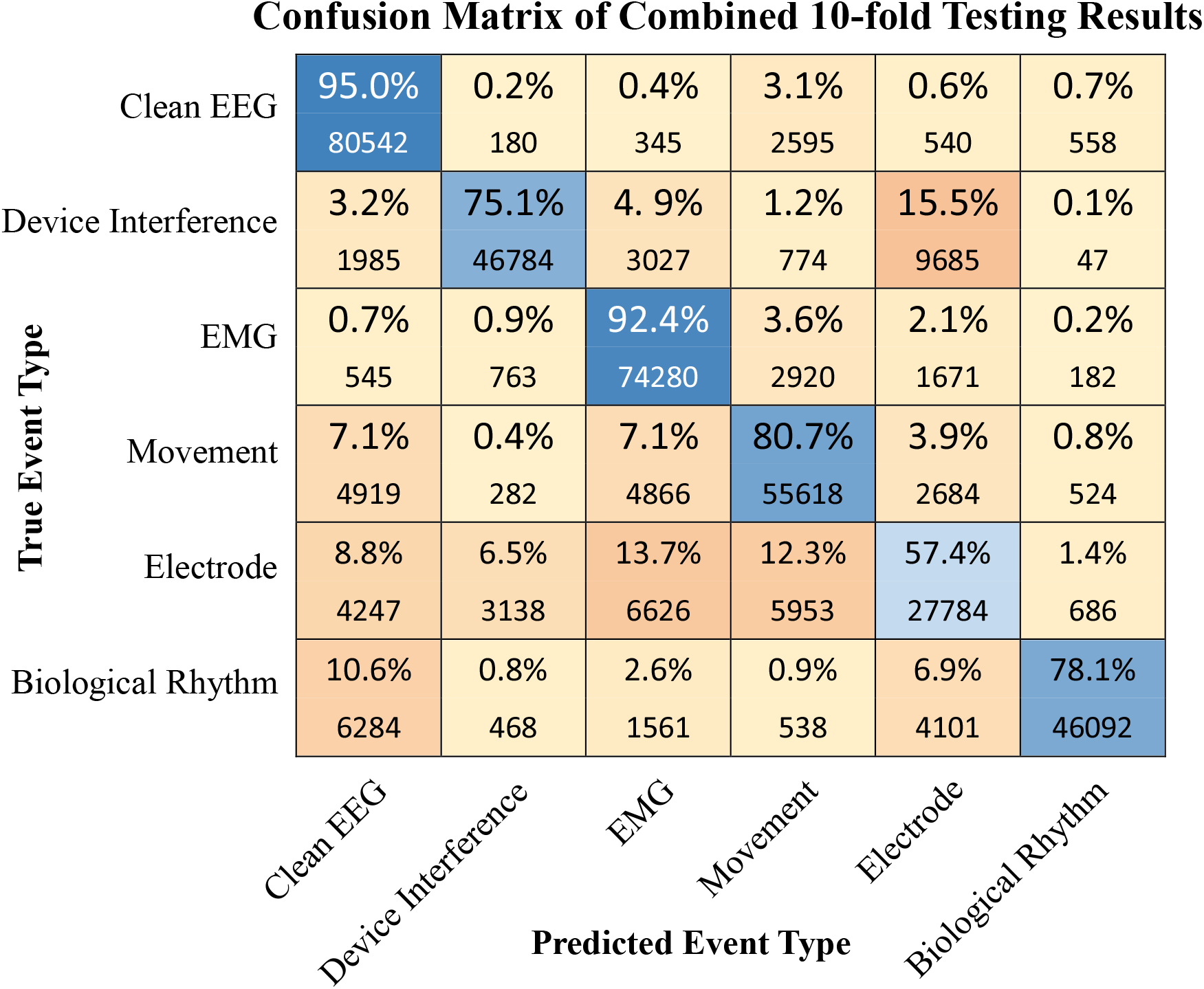
Confusion matrix of event classification on the Balanced Data, summarised over all folds in the ten-fold cross validation. The blue cells along the main diagonal represent correct labelling, with the darkness of the colours representing the size of the percentage in the cell. Percentages are calculated as row percentages, representing the proportion of the true labels in each predicted label. The numbers below the percentages are the number of segments that were annotated by an expert as the row event type, and labelled as the column type by the algorithm.

The separation of each artefact class by the residual neural network is shown in in Figure 6. The *MO* and *EMG* clusters overlap (Figure 6) supporting the finding that *EMG* artefact is most commonly misclassified as *MO* artefact, and vice-versa (Figure 5). Similarly, the *EL* cluster has considerable overlap with the *EMG, MO*, and *DI* cluster, with these classes representing the majority of misclassified *EL* artefacts (Figure 5).

**Figure 6:**
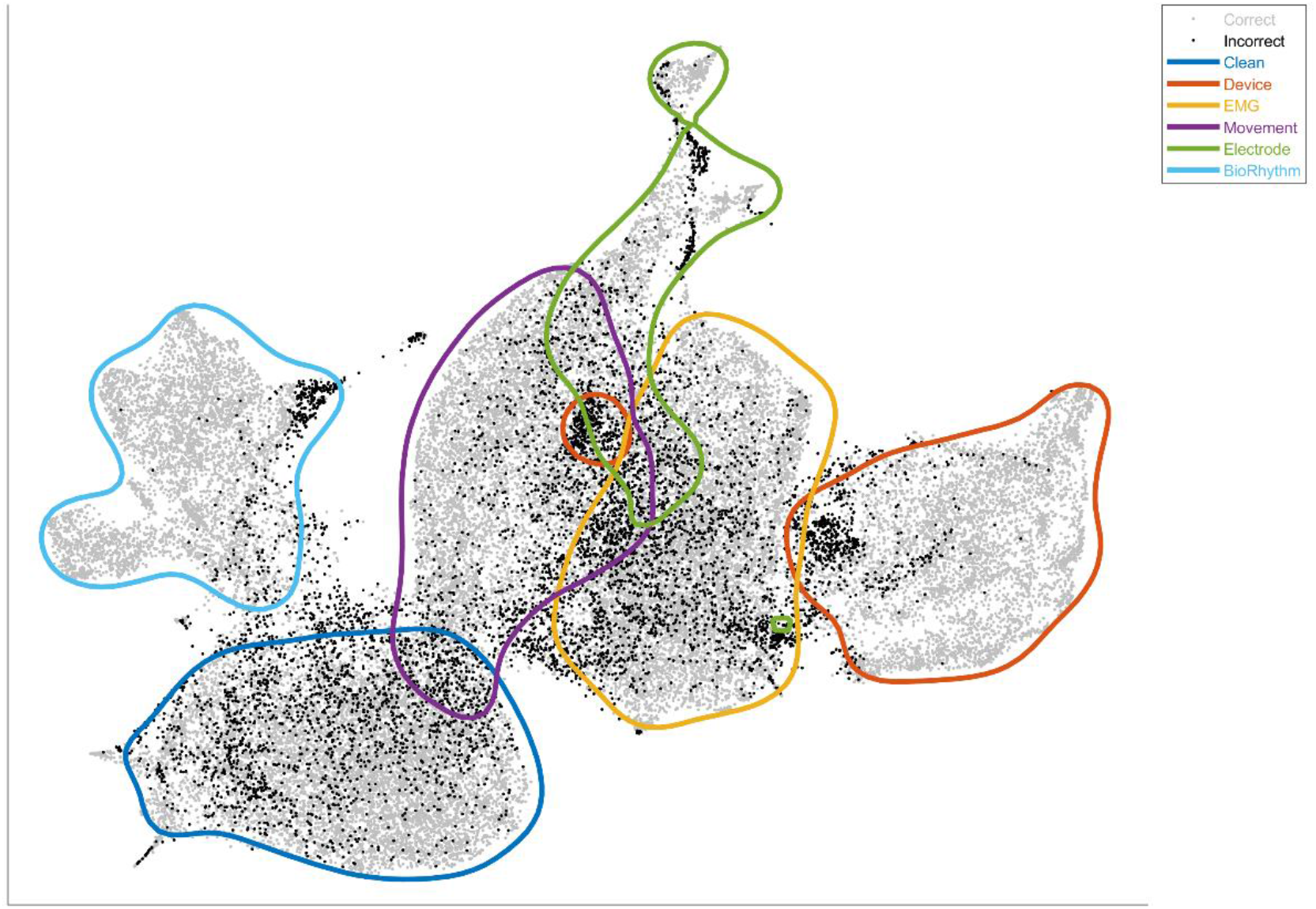
Visualization of the algorithms separation of artefact classes using 10% of the dataset and UMAP dimension reduction. Late stage (fully connected layer) used as an input into the UMAP algorithm. Points shaded grey represent segments correctly identified, and black represents incorrect identification. Coloured contours are used to show the density of annotated artefact classes in the 2 dimensional space.

Applying the artefact detection algorithm to the entire EEG recordings resulted in an overall artefact burden of 30% over all of the channels (sum of five artefact types, Table 3). A seizure detection algorithm was initially used as an additional layer of automated annotation to exclude seizure events [32]. Figure 7 shows the amount of data labelled as *Clean* by the artefact detection algorithm for each channel in the Double Banana bipolar montage, with Table S.5 showing the distribution of events for each channel. The channels at the top of the head (F4-C4, C4-P4, F3-C3, C3-P3, Fz-Cz, Cz-Pz) had a greater proportion of clean data (>75%) than other channels.

**Table 3:**
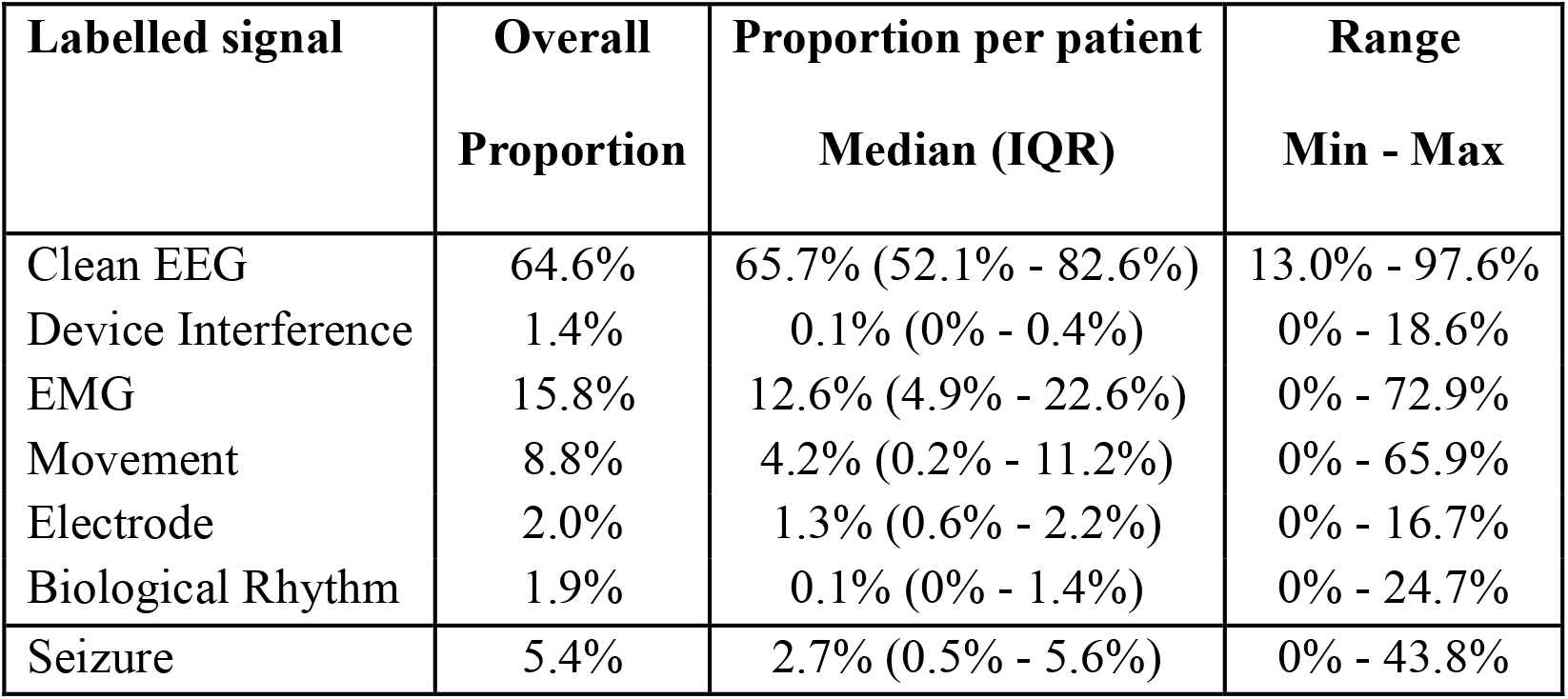
Proportion of each event type as classified by the artefact detection algorithm in complete EEG recordings, after processing by seizure detection.

| Labelled signal | Overall Proportion | Proportion per patient Median (IQR) | Range Min - Max |
| --- | --- | --- | --- |
| Clean EEG | 64.6% | 65.7% (52.1% - 82.6%) | 13.0% - 97.6% |
| Device Interference | 1.4% | 0.1% (0% - 0.4%) | 0% - 18.6% |
| EMG | 15.8% | 12.6% (4.9% - 22.6%) | 0% - 72.9% |
| Movement | 8.8% | 4.2% (0.2% - 11.2%) | 0% - 65.9% |
| Electrode | 2.0% | 1.3% (0.6% - 2.2%) | 0% - 16.7% |
| Biological Rhythm | 1.9% | 0.1% (0% - 1.4%) | 0% - 24.7% |
| Seizure | 5.4% | 2.7% (0.5% - 5.6%) | 0% - 43.8% |

**Figure 7:**
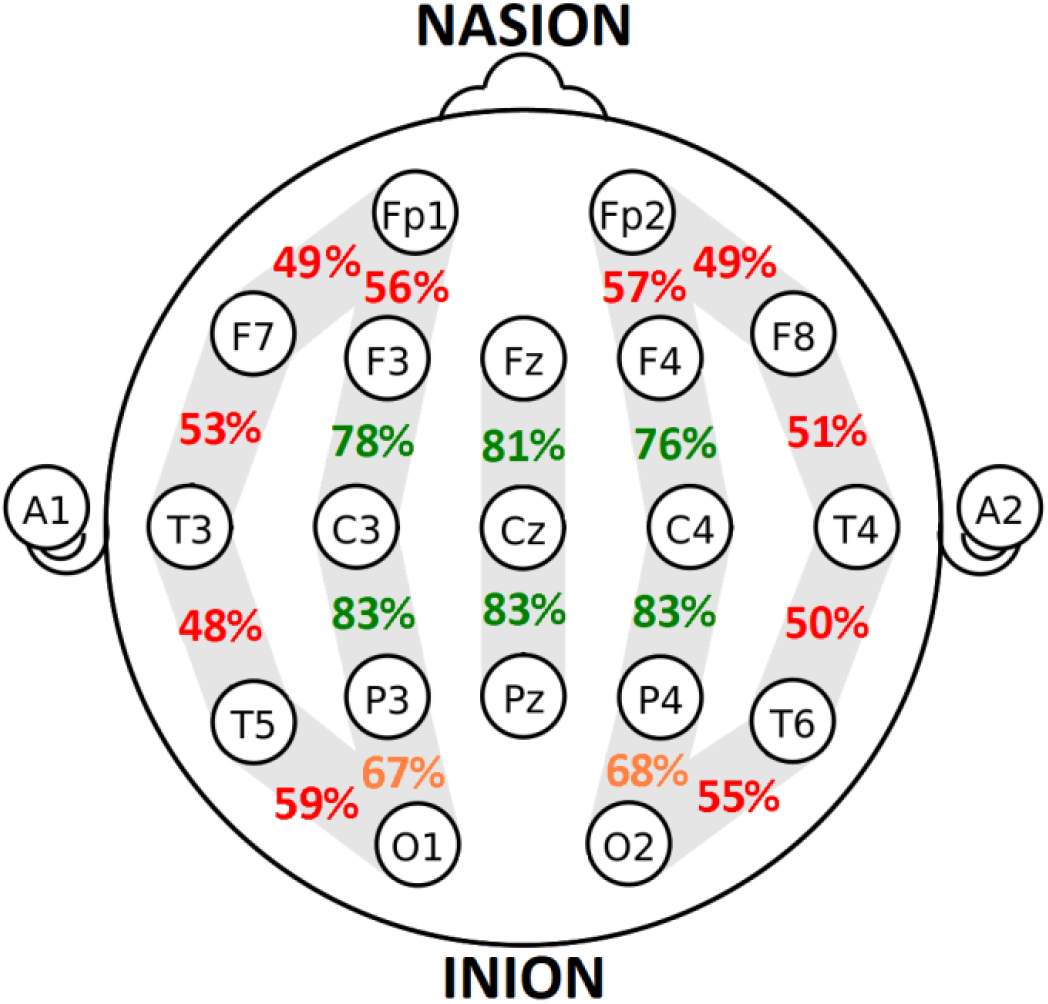
Proportion of clean data in each bipolar channel from the international 10-20 system in the Double Banana montage. Estimated from applying the artefact detection algorithm to the entire training dataset.

### 3.2 Validation Results

The accuracy of our artefact detection algorithm (trained on all of the Balanced Dataset) when applied to the validation dataset is shown in Figure 8. A 6 Class accuracy of 73% was achieved, with a binary accuracy of 87%, sensitivity of 86%, AUC of 0.872 and F1 of 0.899. The accuracy was significantly lower than estimated with 10-fold cross-validation on the developmental set (the 95% bootstrapped confidence intervals for differences in both 6 Class accuracy and binary accuracy did not contain zero, with [2.09, 27 .05] for 6 Class and [2.72 16.22] for binary).

**Figure 8:**
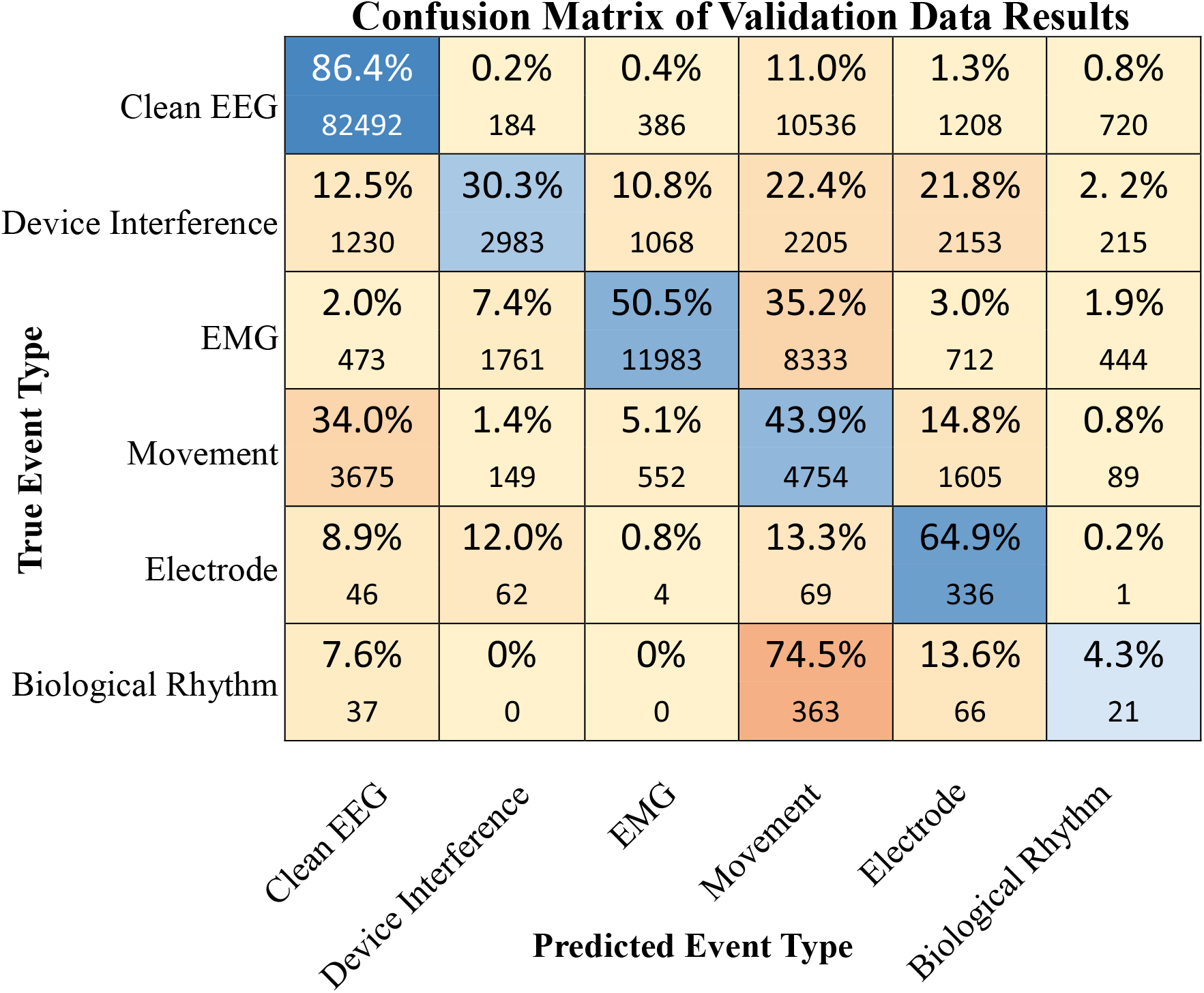
Confusion matrix of event classification on the validation data. The blue cells along the main diagonal represent correct labelling, with the darkness of the colours representing the size of the percentage in the cell. Percentages are calculated as row percentages, representing the proportion of the true labels in each predicted label. The numbers below the percentages are the number of segments that were annotated by an expert as the row event type, and labelled as the column type by the algorithm.

A potential explanation for the reduction in algorithm performance was the differences in process of annotation between the training and validation datasets. For example, the median average duration for each event type annotated was longer in the validation dataset, with median duration of 38 s for *EMG*, 28 s for *MO* compared to 15 s and 12 s in the training dataset (Table S.1). The annotated segments from the training data also had different EEG characteristics compared to the annotated segments in the validation dataset (Figure 9). For a number of the artefact classes in the validation data, the mean frequency is closer to that of the clean segments in the training dataset. In the majority of the classes, including *Clean*, the mean amplitude is increased in the validation data.

**Figure 9:**
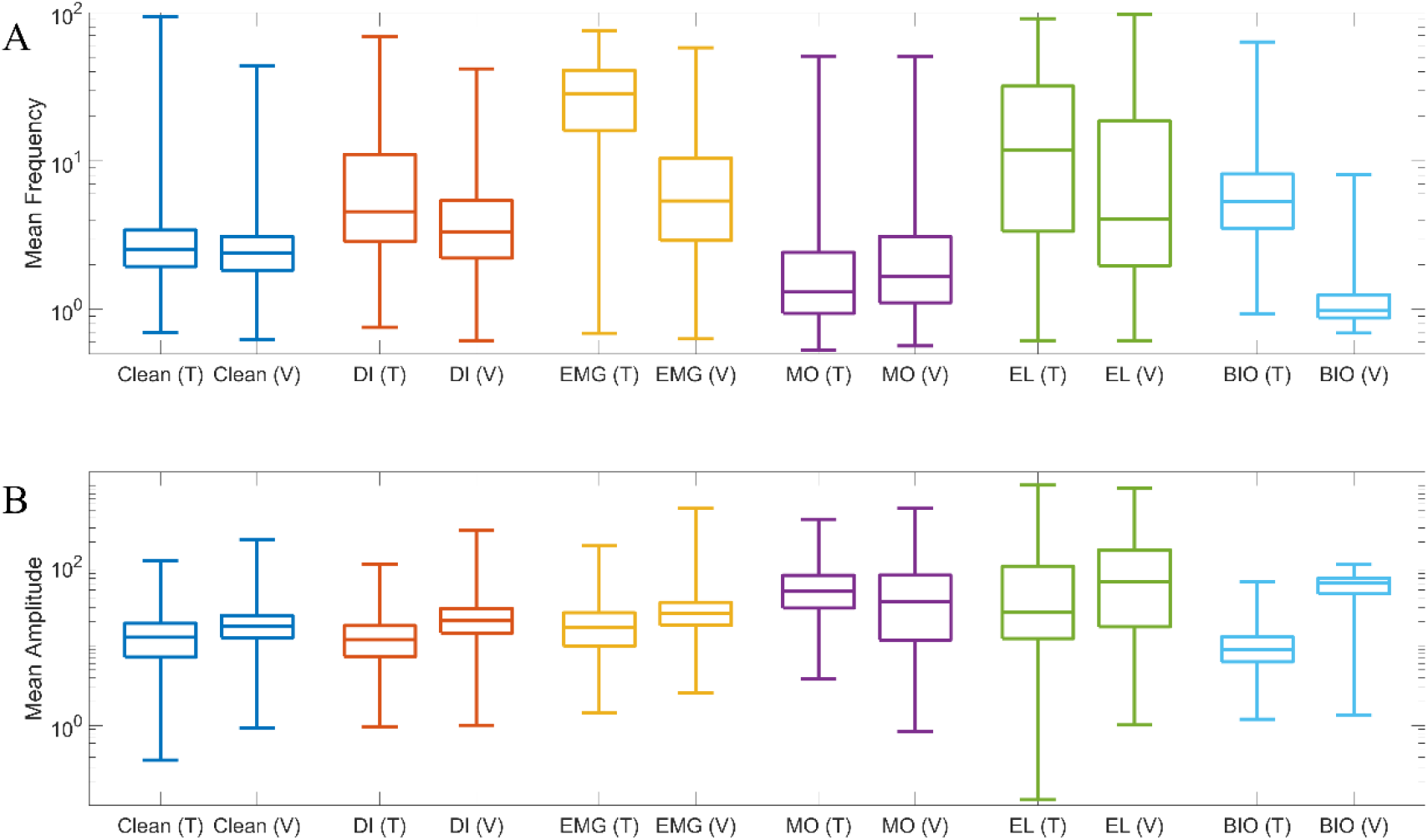
A comparison of A) the Mean Frequency of the EEG, and B) the Mean Amplitude of the EEG in the annotated segments between training and validation datasets. One tenth of the training data compared to all of the validation annotations. Distributions of *Clean* segments from the training dataset is only one tenth of dataset. Training (T) and Validation (V) datasets for the *Clean* segments, and artefact segments *Device Interference* (DI), *Electromyography* (EMG), *Movement* (MO), *Electrode* (EL), and *Biological Rhythm* (BIO). A) *Clean*, DI, MO and EL segments are similar in Mean Frequency in both training and validation, with substantial differences seen in EMG and BIO segments. B) The distribution of Mean Amplitude differs largely for BIO, as well as other differences in centre for DI,

## 4. Discussion

We have developed an algorithm for detecting several types of common artefacts in the neonatal EEG. This algorithm uses a deep residual net trained on EEG data annotated by a human expert. The main innovation of this paper lies in the development of a unique system of artefact detection that not only detects the presence of artefact in neonatal EEG but also classifies the type of artefact. This algorithm can be used i) to aid bedside nursing staff in monitoring EEG quality in real time, in order to take corrective actions (e.g. detection of *EL* artefact can suggest poor contact of specific electrodes), and ii) to support EEG review by clinician’s and complement future diagnostic tools, such as seizure detectors, lesion detectors, or EEG background classifiers [28, 32-35].

The detection algorithm differentiated between artefact and artefact-free EEG with an accuracy of 95% and identified the type of artefact with an accuracy of 85% on a developmental subset of 79 EEG recordings. Applying the artefact detection algorithm to the entire EEG dataset showed that the commonly used electrode positions used in long-term, limited channel monitoring (frontal, central, and parietal locations) were the least susceptible to artefacts. The detection accuracy of the algorithm on an independent validation set was 87% (temporal averaging was not applied in this case), suggesting practical utility.

The accuracy of the neural net stage of the algorithm to detect artefact in general was high (95% accuracy, Table 2 and Figure 5), although the performance of the neural net in terms of identifying specific artefacts varied with the type of artefact to be detected (from 57% to 92%, Figure 5). Artefacts that were the most difficult to detect were those that were i) rarer in the dataset such as *DI* and *BIO* (Table S.1), ii) more variable with several distinct phenomena represented with a class such as *BIO* and *EL*, iii) contained periods of uncontaminated EEG activity such as *EL* and *BIO*, or iv) contained a mixture of artefact types such as *EMG* and *MO* (Figure 4B). These deficits may be overcome by improving the classifier design and implementation, improving the time resolution of the annotation and redefining the artefact classes, e.g. either combining movement and EMG or adding a separate EMG + movement class.

The accuracy of the artefact detection algorithm was significantly reduced in the validation data compared to results obtained using cross-validation on the developmental set. Inspection of the quantitative summary measures between correct and incorrect detections (Figure 9) suggests that portions of the validation data within several artefact classes have different EEG characteristics to those present in the training/development dataset. While both datasets were obtained from the same hospital and using the same EEG machine, there are several technical and practical differences between datasets: EEG acquisition, annotation, and lastly the generalizability of the training data. We have shown that the 4-channel montage used in the validation dataset is particularly robust to the presence of artefact, which i) reduces the probability of artefact (Figure 7) and ii) potentially dilutes the manifestation of artefact on the EEG. The annotation of artefact is also not trivial and susceptible to different interpretations (subjective). The reliance on human annotations is the main challenge and limitation for developing computational artefact detectors/classifiers. There are no objective criteria for the minimal presence (i.e., detection threshold) of artefacts, and it is also well known that several artefacts are often seen at the same time (e.g. *MO* and *EMG*). More annotated recordings using a consistent process of annotations from multiple centers would improve the performance and generalisability of the algorithm.

There is a dearth of methods available for artefact detection in the neonatal EEG. Neonatal EEG is different from the EEG of children and adults, it has significant power in lower frequencies unique patterns relating to normal and abnormal function and can provide useful information to clinicians when recorded with limited channel montages [6, 36]. These differences mean that artefact detection methods based on the analysis of adult EEG (a more mature field), at best, need to be re-trained on neonatal training data and, at worst, cannot be effectively implemented. Nevertheless, there have been attempts at detecting artefact in neonatal EEG. Schetinin and Schult [20] use a ‘bag of features’ classified by a hybrid decision tree/polynomial neural network to detect artefacts during sleep in the neonatal EEG with a binary accuracy of 73.5±2.8%. In this paper, we achieve comparable accuracy (median binary accuracy 92%, IQR: 86-95%) using a similar process (bag of features classified by a support vector machine) based on our earlier work [5]. We show that the use of neural networks significantly improves the accuracy beyond these methods (median binary accuracy 98%, IQR: 95-99%) to a level that is comparable to the state of the art in children and adults [37].

We have developed a neonatal EEG artefact detection algorithm based on a residual neural network. The algorithm achieves good accuracy for both identifying the presence and type of artefact contaminating an EEG recording. Future work includes: i) the investigation of different neural network architectures, data augmentations, training options and data processing, ii) developing potential improvements to the process of annotating EEG artefacts, iii) performing studies of inter-rater agreement to set appropriate performance benchmarks, and iv) the acquisition of larger, multi-center datasets to allow algorithms to experience a large diversity of artefacts and compensate for potential sources of heterogeneity.

## Supplementary Material

**Table S.1:** Identified events annotated in the 79 EEG recordings

| Labelled signal | #Events (%) | Average Duration (s),<br>median (range) | Total Duration of events<br>(min) |
| --- | --- | --- | --- |
| Clean EEG | 359 (11.0) | 127 (4 – 1682) | 1250.2 |
| Device Interference | 232 (7.1) | 12 (2 – 450) | 75.8 |
| EMG | 1167 (35.7) | 15 (2 – 3135) | 881.4 |
| Movement | 540 (16.5) | 12 (1 – 315) | 181.2 |
| Electrode | 894 (27.4) | 4 (1 – 520) | 144.1 |
| Biological Rhythm | 78 (2.4) | 40 (8 – 2384) | 134.4 |
| Total | 3270 |  | 2667.1 (44.5 h) |

**Table S.2:** Number of segments in each fold of the cross-validation

| Fold | Number of segments in training (Balanced Data) |  |  |  |  |  |  |
| --- | --- | --- | --- | --- | --- | --- | --- |
|  | Total | Clean | Device<br>Interference | EMG | Movement | Electrode | Biological<br>Rhythm |
| 1 | 33613 | 7743 | 599 | 13090 | 1117 | 903 | 10161 |
| 2 | 32584 | 9179 | 208 | 5528 | 13179 | 3605 | 885 |
| 3 | 60476 | 10988 | 15991 | 5409 | 146350 | 747 | 12706 |
| 4 | 29350 | 9363 | 6947 | 7782 | 3531 | 502 | 1225 |
| 5 | 37907 | 5817 | 2122 | 14509 | 6617 | 8047 | 795 |
| 6 | 40864 | 8656 | 99 | 8526 | 11133 | 10147 | 2303 |
| 7 | 39638 | 10852 | 4528 | 8225 | 8646 | 4680 | 2527 |
| 8 | 47496 | 8977 | 1746 | 6029 | 1051 | 14825 | 14868 |
| 9 | 24915 | 6490 | 4730 | 5170 | 5424 | 2261 | 840 |
| 10 | 56951 | 6695 | 25332 | 6093 | 3560 | 2537 | 12734 |

**Table S.3:**
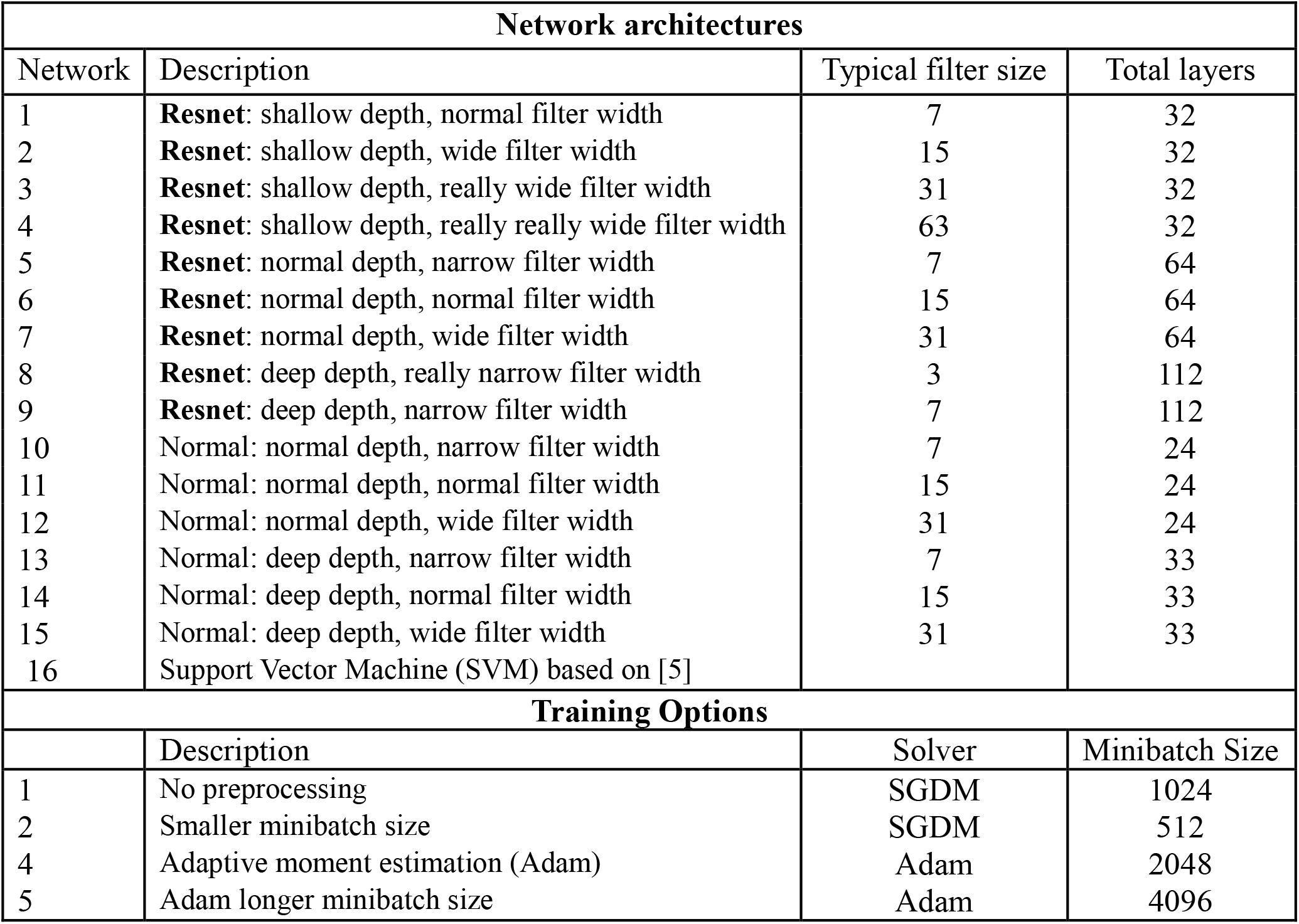
Neural Network architectures and training options considered.

**Table S.4:**
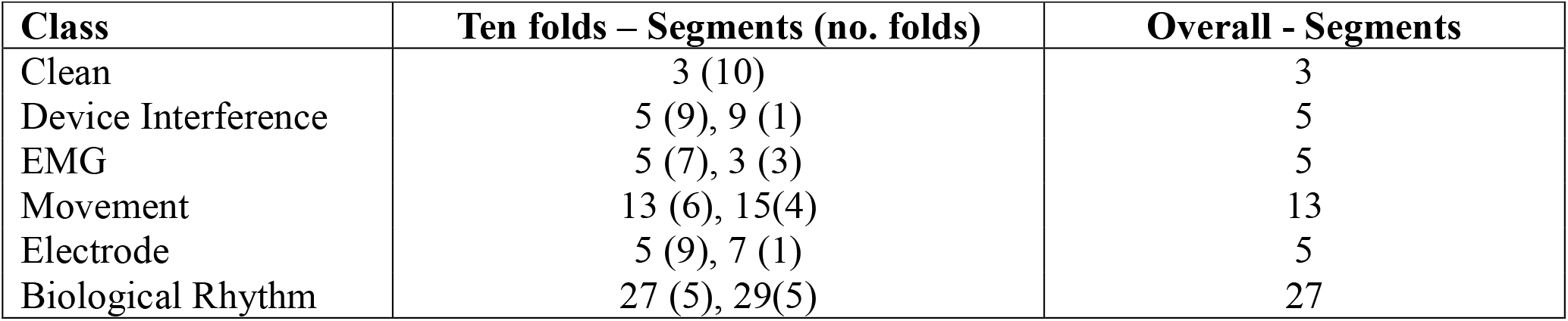
Optimal number of segments in smoothing window.

| Class | Ten folds – Segments (no. folds) | Overall - Segments |
| --- | --- | --- |
| Clean | 3 (10) | 3 |
| Device Interference | 5 (9), 9 (1) | 5 |
| EMG | 5 (7), 3 (3) | 5 |
| Movement | 13 (6), 15(4) | 13 |
| Electrode | 5 (9), 7 (1) | 5 |
| Biological Rhythm | 27 (5), 29(5) | 27 |

**Table S.5:** Distribution of label given by artefact detection algorithm or seizure detection algorithm when applied to whole recordings (n=79). Bolded percentages in the Clean column are above 70%, and in the artefact columns are above 20%.

| Channel | Event type labelled by artefact detection algorithm, or seizure detection algorithm (%) |  |  |  |  |  |  |
| --- | --- | --- | --- | --- | --- | --- | --- |
|  | Clean | Device Interference | EMG | Movement | Electrode | Biological Rhythm | Seizure |
| Fp2-F4 | 57.0 | 1.0 | **28.9** | 6.2 | 2.6 | 0.6 | 3.7 |
| F4-C4 | **78.6** | 1.4 | 6.3 | 7.4 | 0.9 | 0.3 | 5.1 |
| C4-P4 | **84.2** | 0.9 | 0.6 | 7.5 | 0.7 | 1.0 | 5.2 |
| P4-O2 | 68.3 | 1.5 | 4.2 | 11.7 | 1.5 | 7.2 | 5.6 |
| Fp1-F3 | 58.2 | 0.5 | **29.4** | 5.8 | 2.2 | 0.1 | 3.8 |
| F3-C3 | **76.6** | 1.5 | 7.9 | 7.7 | 1.5 | 0.1 | 4.8 |
| C3-P3 | **84.5** | 1.8 | 0.1 | 6.9 | 0.7 | 0.0 | 6.0 |
| P3-O1 | **69.4** | 1.3 | 4.4 | 13.3 | 1.0 | 4.1 | 6.6 |
| Fp2-F8 | 49.6 | 1.7 | **31.6** | 7.0 | 3.3 | 0.6 | 6.2 |
| F8-T4 | 53.5 | 1.6 | **25.4** | 9.9 | 3.2 | 1.1 | 5.4 |
| T4-T6 | 48.7 | 1.5 | **26.8** | 11.1 | 3.3 | 3.4 | 5.1 |
| T6-O2 | 59.8 | 1.8 | 16.5 | 11.4 | 2.7 | 2.7 | 5.1 |
| Fp1-F7 | 50.0 | 1.8 | **32.8** | 6.9 | 2.6 | 0.5 | 5.3 |
| F7-T3 | 51.2 | 1.8 | **28.6** | 9.5 | 2.8 | 1.1 | 5.0 |
| T3-T5 | 50.6 | 1.3 | **25.8** | 12.3 | 2.6 | 2.5 | 4.8 |
| T5-O1 | 56.1 | 1.0 | 15.3 | 11.8 | 2.1 | 7.8 | 5.9 |
| Fz-Cz | **82.4** | 2.5 | 0.0 | 5.3 | 0.9 | 1.7 | 7.3 |
| Cz-Pz | **84.5** | 1.0 | 0.0 | 7.2 | 0.6 | 0.0 | 6.7 |

**Figure S.1:**
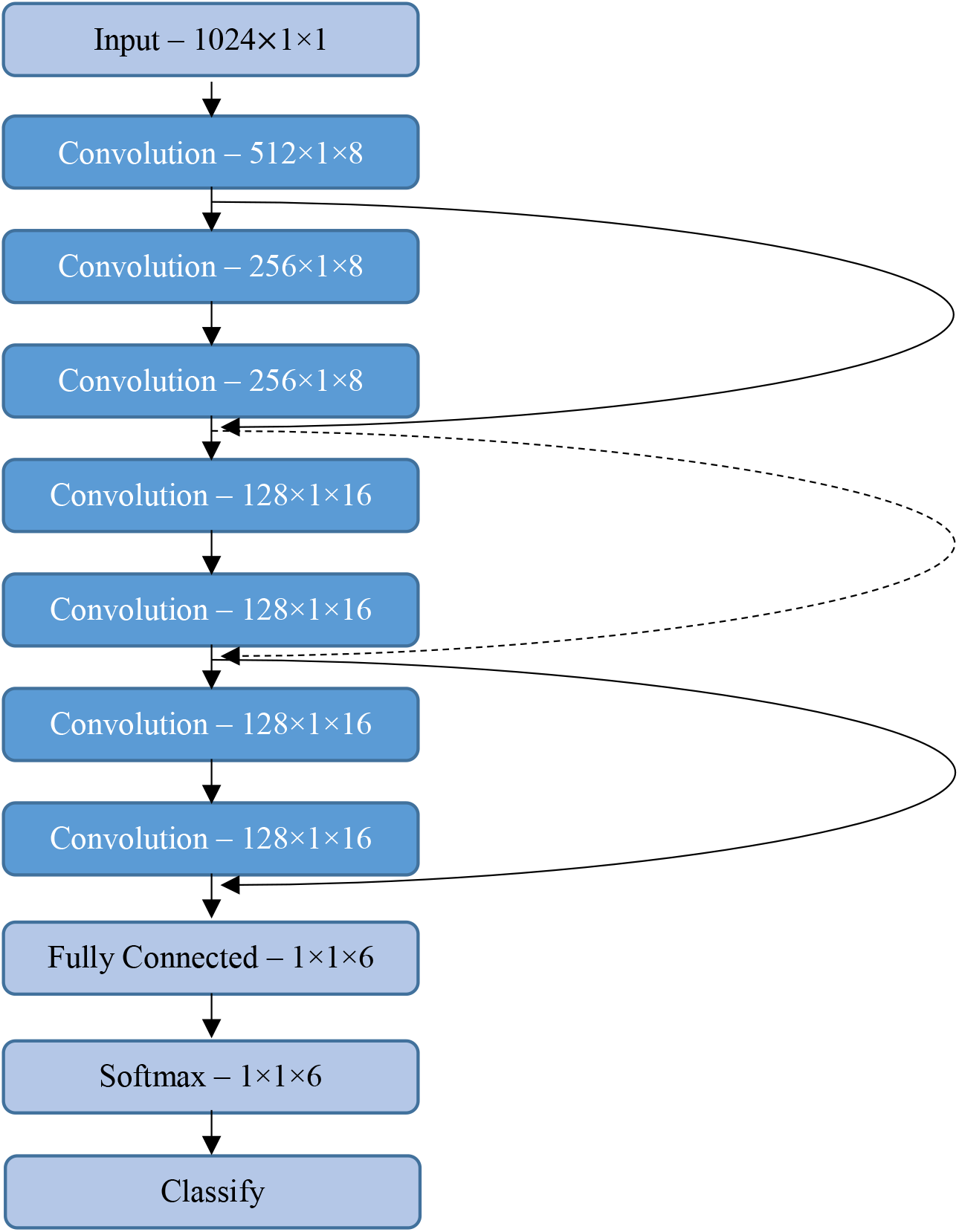
The Deep Residual CNN used in the artefact detection system. Curved lines are shortcuts. Dashed shortcuts increase dimensions with padding.

**Figure S.2:**
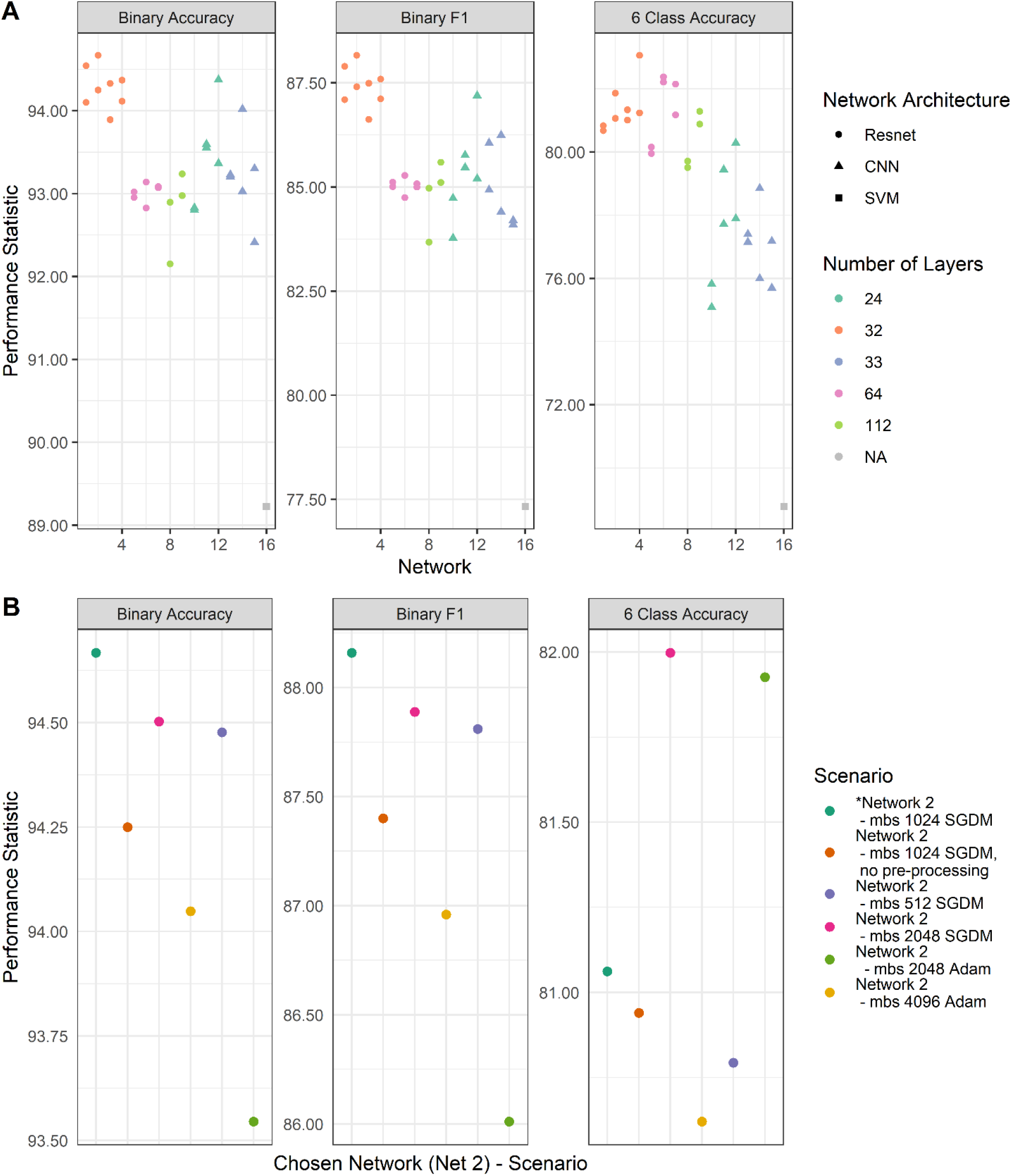
Performance of the trialled neural network architectures, and training options of the best performing network. A) Each networks except the SVM was trained twice, each time with different starting seed. B) The far-left network (*) is Network 2 from A. Scenario labels; SGDM is stochastic gradient descent method, Adam is the Adam method, and mbs is minibatch size.

## Acknowledgments

This work was supported by the National Health and Medical Research Council of Australia (JAR, SV, APP1144936), the Rebecca L. Cooper Foundation (JAR, PG2018109), EU Marie Skłodowska Action (SV, H2020-MCSA-IF-656131), Academy of Finland (SV, #335778, #314450, #332017), Helsinki University Hospital, Lastentautien tutkimussaatio, Aivosaatio, and Sigrid Juselius Foundation.

